# A Multi-Institution Biobanking Pipeline for Primary Human Satellite Cells and Fibro-Adipogenic Progenitors

**DOI:** 10.64898/2026.07.15.738758

**Authors:** Frank S. Pittman, Adam Rauff, Grace E. Privett, Alis Balayan, Severin Ruoss, Robert E. Guldberg, Catherine M. Robertson, Adam J. Engler, Samuel R. Ward, Nick J. Willett

**Affiliations:** Department of Bioengineering, Phil and Penny Knight Campus for Accelerating Scientific Impact, University of Oregon; Eugene, Oregon, United States; Biomedical Sciences Program, UC San Diego, La Jolla, California, United States; Department of Orthopaedic Surgery, UC San Diego, La Jolla, California, United States; Chien-Lay Department of Bioengineering, UC San Diego, La Jolla, California, United States; Sanford Consortium for Regenerative Medicine, La Jolla, California, United States; Department of Radiology, UC San Diego, La Jolla, California, United States; Department of Biomedical Engineering, Oregon Health & Science University; Portland Oregon, United States; Department of Orthopaedics & Rehabilitation, Oregon Health & Science University; Portland, Oregon, United States; The Veterans Affairs Portland Health Care System, Portland, Oregon, United States

**Keywords:** Muscle, biobanking, satellite cells (SCs), fibro-adipogenic progenitors (FAP)

## Abstract

Satellite Cells (SCs) and Fibro-Adipogenic Progenitors (FAPs) are muscle-resident cell populations crucial for maintaining skeletal muscle homeostasis and coordinating regeneration after injuries. However, primary human SCs and FAPs are difficult to co-isolate, and their broad use in translational research has been limited by a lack of standardized biobanking protocols. Recently, we published a protocol for efficient co-isolation of SCs and FAPs from human skeletal muscle. Here, we extend those efforts to establish a comprehensive pipeline for the cryopreservation, cold-chain transport, and independent-site utilization of human SCs and FAPs. Cells taken through this pipeline maintained lineage-specific markers, including Pax7, MyoD and CD56 for SCs, and PDGFRα and TE7 for FAPs, indicating retention of their pre-biobanking phenotype. Furthermore, SCs demonstrate robust myogenic differentiation capacity, and FAPs demonstrate both fibrogenic and adipogenic differentiation capacity post-transport. Finally, previously biobanked SCs were incorporated into *in vitro* 3D muscle constructs, demonstrating their utility for human-based New Approach Methodologies (NAMs). This framework for multi-site collaboration facilitates broader access to human primary muscle cells, which will improve the scalability and translatability of human-based NAMs for skeletal muscle research.

## Introduction

Skeletal muscle is a metabolically and mechanically active tissue that is crucial for everyday functions including locomotion, respiration, thermogenesis, and glucose metabolism^1,2^. Severe skeletal muscle injury, such as volumetric muscle loss, can result in the replacement of healthy muscle with non-contractile fibrotic and fatty tissue, thereby impairing muscle function and reducing quality of life^3,4^. Two critical cell populations involved in muscle regeneration are satellite cells (SCs) and fibro-adipogenic progenitors (FAPs). SCs, the predominant myogenic stem cell type, reside between the basal lamina and the muscle fiber in a quiescent state. Upon muscle injury, SCs become activated, subsequently proliferating, differentiating, and fusing to form new muscle fibers^5,6^. FAPs are muscle-resident mesenchymal cells that synthesize and remodel the muscle extracellular matrix. Furthermore, FAPs support myogenesis via paracrine signaling with SCs^7^. In successful regeneration following muscle injury, FAPs are cleared from the regenerative niche via TNFα-mediated apoptosis^8,9^. However, unsuccessful regeneration, such as following volumetric muscle loss, leads to disruption of FAP-SC crosstalk and pathological differentiation of FAPs into myofibroblasts (fibrogenic differentiation) and adipocytes (adipogenic differentiation)^10–12^. This dysregulation of FAPs differentiation results in fibrotic and fatty infiltration of the skeletal muscle. Given their indispensable role in muscle regeneration, and the effect of their dysregulation on skeletal muscle composition, SCs and FAPs are promising therapeutic targets.

Animal models have provided foundational knowledge of the mechanisms driving pathological regeneration following severe muscle injury^13^. However, current clinical treatments derived from preclinical studies cannot reliably restore muscle mass or function^14–16^. To close this translational gap, human-based models are critical to advance our understanding of human-specific mechanisms of muscle pathologies. The development of human-based New Approach Methodologies (NAMs) is a major priority, yet broad access to validated populations of primary human muscle-derived cells is required to do so.

While considerable progress has been made to develop isolation and culture protocols for human FAPs and SCs^17–19^, many gaps limit accessibility to these cells for multi-institutional research. Cell yield from human muscle tissue samples is typically low, and the ethical, regulatory, and logistical barriers to acquiring human tissue samples can be substantial^20,21^. These barriers are especially relevant for institutions without access to surgical centers. Compounding these constraints, human-specific SC and FAP isolation markers are not standardized across the field. While CD56 and PDGFRα are widely used to identify SCs and FAPs, respectively, the broader panel of markers to improve purity or exclude contaminating populations varies across published protocols^12,17,22–24^. Additionally, many commercial vendors for primary human muscle cells do not publish their isolation methods. Thus, cell populations that are isolated by independent research groups, or supplied by commercial biobanks, can differ in ontological identity. Critically, while some commercial biobanks separately offer human SCs, myoblasts, or FAPs, no standardized biobanking infrastructure has been reported for matched pairs of Pax7^+^ SCs and PDGFRα^+^ FAPs co-isolated from the same donor. Addressing these gaps will dramatically enhance and accelerate the development of human-specific skeletal muscle NAMs.

The present study directly addresses these gaps in skeletal muscle regenerative research. Building off previously published methods for co-isolation of human skeletal muscle SCs and FAPs ^17^, we report detailed methods for an end-to-end pipeline to biobank, ship, and utilize these cell populations. This framework encompasses co-isolation from human patients^17^, cryopreservation and cold-chain transport, and verification of post-thaw lineage and functional differentiation capacity. Additionally, we utilize derived SCs to successfully generate 3D tissue engineered muscle organoids, demonstrating their potential for use in NAMs. This framework for multi-site collaboration facilitates broader access to reliable primary muscle cells, which will enhance the development of reproducible and clinically translatable human-based NAMs for skeletal muscle research.

## Methods

### Overview of Biobanking Pipeline

Use of human tissue followed the principles outlined in the Declaration of Helsinki. The experimental protocol was approved by UC San Diego’s Institution Review Board protocol 181569 and written informed consent was obtained from participating subjects. A graphical representation of the established biobanking pipeline is shown in Figure 1. The sample collection, SC and FAP co-isolation, and assessment of pre-freeze multi-lineage differentiation capacity were reported by Balayan et al.^17^. In the present study, cells were cryopreserved and stored in the Wu Tsai Human Performance Alliance Biobank (https://humanperformance.ucsd.edu/?page_id=460) ^25^. Cells were then shipped via cold chain to an independent research site (University of Oregon), where they were stored in the liquid nitrogen vapor phase until use. During initial expansion post-thaw, lineage marker expression of SCs and FAPs was verified by immunofluorescence (IF) imaging and flow cytometry. Then, the functional differentiation capacity of SCs into myotubes and FAPs into adipocytes and myofibroblasts was assessed via IF imaging. Lastly, utilization of SCs was demonstrated by generating *in vitro* tissue-engineered skeletal muscle constructs.

**Figure 1:**
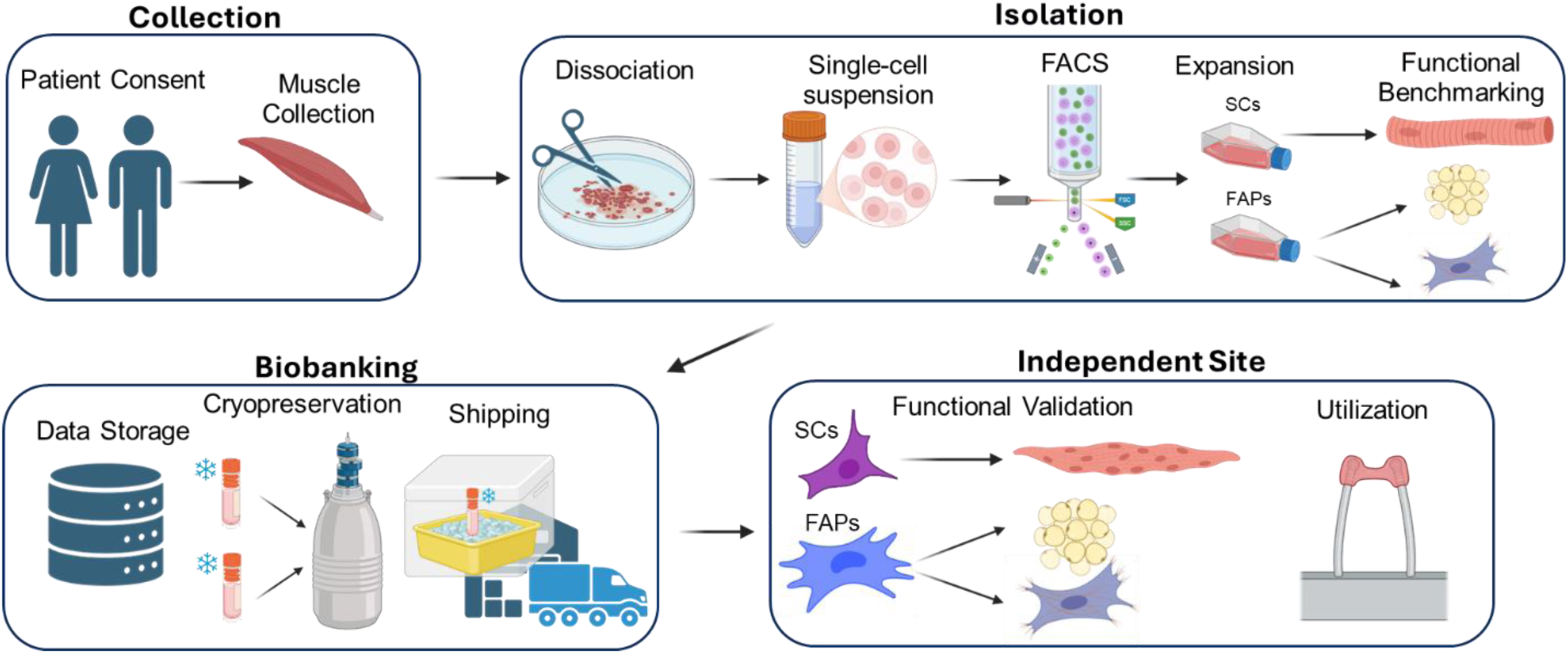
Overview of biobanking pipeline. Muscle tissue specimen collection and SC/FAP co-isolation were previously described by Balayan et al.^17^. Cataloguing of de-identified patient demographic data, cryopreservation and cold-chain shipping of SC and FAP samples were performed by the Wu-Tsai Biobank at UC San Diego. Confirmation of SC and FAP lineage marker expression and functional differentiation were performed after receipt at the University of Oregon. SCs were then utilized in a 3D tissue-engineered muscle construct. Created in BioRender. Willett, N. (2026) https://BioRender.com/hupwot8.

### Isolation of SCs and FAPs

Satellite cells (SCs) and fibroadipogenic progenitors (FAPs) were isolated from the gracilis of one donor (female, age 25) according to a previously published protocol^17^. Briefly, skeletal muscle samples were minced and enzymatically digested using a combination of Dispase II (Gibco, 2 U/mL) and collagenase, type 1 and 2 (Worthington Biochemical, 50 U/mL) for 90 minutes at 37°C with constant agitation. After tissue digestion, the cells were strained (70 µm, Fisher Scientific) and red blood cells were lysed using Ammonium-Chloride-Potassium (ACK, Quality Biological). For fluorescence-activated cell sorting (FACS), cells were then stained with fluorophore-conjugated primary antibodies: anti-CD45-FITC (BD Bioscience), anti-CD31-APC-CY7 (1:40, BD Bioscience), anti-CD56-PE-CF594 (1:40, BD Bioscience) and eFluor 506 Viability dye (1:100, eBioscience). FACS with Aria Fusion or Aria II (BD Bioscience) was used to sort CD56+CD31-CD45-(SCs) and CD56-CD31-CD45-(FAPs) cells.

### Fresh Cell Culture and Cryopreservation

Populations of sorted SCs and FAPs were expanded in proliferation media, consisting of Dulbecco’s Modified Eagle’s Medium (DMEM) low glucose (Gibco) supplemented with 20% fetal bovine serum (FBS, Omega Scientific), 5 ng/mL recombinant human fibroblast growth factor 2 (rhFGF2, R&D Systems) and 1% penicillin/streptomycin (P/S, Gibco). Proliferation media was replaced every 2–3 days during expansion. To passage SCs and FAPs, the cells were washed with Dulbecco’s Phosphate Buffered Saline (DPBS, Gibco) and lifted using 0.25% Trypsin-Ethylenediaminetetraacetic acid (EDTA, Gibco) for 5 minutes at 37°C. Trypsin-EDTA was neutralized using an equal volume of proliferation media and the cells were pelleted at 400*g* for 5 minutes. The cells were then sub-cultured at 3500 cells/cm^2^. SCs and FAPs were cryopreserved at passages 3-5 using Cryostor CS10 (Stem Cell Technologies) at 1 million cells per 1ml of Cryostor per vial. The cells were placed in a Mr. Frosty freezing container (Nalgene) to allow −1°C/minute cooling rate in −80°C. The following day, the frozen cells were transferred to liquid nitrogen for long-term storage. The cryopreserved SCs and FAPs are available via the Wu Tsai Human Performance Alliance biobank (<u>humanperformance.ucsd.edu</u>) upon reasonable request.

### Sample Shipping

On shipping day, samples were moved from the Biobank’s liquid nitrogen storage into a liquid nitrogen dry vapor shipping container. Samples were shipped overnight by Cryoport Systems LLC. The shipment was coordinated to be during two consecutive working days without a holiday. The recipient site received tracking information and access to Cryoport’s proprietary monitoring dashboard. During shipment, average internal temperature was −194.2 °C (minimum: −209.3 °C, maximum: −188.6 °C) and average external temperature was 20 °C (minimum: 13.3 °C, maximum: 27.0 °C). Average humidity was 52% (minimum: 40%, maximum: 64%) and average pressure was 14.5 PSI (minimum: 11.8 PSI, maximum: 14.8 PSI). Upon arrival, samples were immediately transferred to a liquid nitrogen dewar for storage in the vapor phase until use.

### SC and FAP Expansion

At the recipient site, SCs and FAPs were expanded following Supplemental Protocols A, B, and C. Samples (passage 4) were removed from frozen storage and swirled in a 37°C water bath for 1 minute to thaw. Cells were then quickly transferred to prewarmed proliferation media prepared as described above. The cell suspension was pelleted at 400g for 5 minutes and cells were counted on a Countess 3 automated cell counter (Invitrogen). All cells (initially stored at 1 million/vial) were re-plated in T175 plastic culture flasks (Thermo Fisher Scientific) in proliferation media. Media was replaced 24 hours after initial plating and subsequently every 2-3 days during expansion. FAPs were expanded to 80-85% confluency and SCs were expanded to 70% confluency before passaging. Once the target confluency was reached, cells were lifted using TrypLE Select Enzyme (Thermo Fisher Scientific) and sub-cultured for use in multilineage differentiation experiments. Any cells not needed for differentiation experiments were cryopreserved for potential future use (Supplemental Protocol D).

### Multilineage Differentiation

Myogenic differentiation was performed using Supplemental Protocol E. Briefly, SCs were seeded at 10,000 cells/cm^2^ and cultured in proliferation media until cells were 100% confluent. Media was then changed to myogenic differentiation media containing DMEM low glucose (Gibco), 5% horse serum (Omega Scientific), 10µg/mL insulin (Sartorius), and 1% P/S (Gibco). Myogenic differentiation media was replaced every 2 days. On day 4 of differentiation, cells were fixed with 4% paraformaldehyde (PFA) for immunofluorescence staining and imaging.

Fibrogenic differentiation was performed using Supplemental Protocol F. FAPs were seeded at 3,000 cells/cm^2^ and cultured in proliferation media overnight. The next day, media was changed to fibrogenic differentiation media containing DMEM low glucose (Gibco), 20% FBS (Gibco), 1% P/S (Gibco), and 10 ng/mL recombinant human transforming growth factor-β1 (TGFβ1, R&D Systems). Fibrogenic differentiation media was replenished every day. On day 4 of differentiation, FAPs were fixed with 4% PFA for immunofluorescence staining and imaging.

Adipogenic differentiation was performed using Supplemental Protocol G. FAPs were seeded at 10,000 cells/cm^2^ and cultured in proliferation media for 4 days. Media was then replaced with StemPro Adipogenesis Differentiation media (Gibco). Adipogenic differentiation media was replenished every 3-4 days. After 21 days of differentiation, cells were fixed with 4% PFA for immunofluorescence or Oil Red O (Milipore Sigma) staining and imaging.

### Immunofluorescence Staining

Immunofluorescence staining was performed using Supplemental Protocol H, which was adapted from previously established protocols^17^. Cells were fixed with 4% PFA for a minimum of 10 minutes at room temperature, washed twice with DPBS for 5 minutes each, permeabilized with 0.2% Triton X (Fisher Scientific) for 10 minutes at room temperature, and washed two times with 2% bovine serum albumin (BSA, Millipore Sigma) for 5 minutes each. Nonspecific binding was blocked with 2% BSA for 1 hour at room temperature followed by overnight incubation with primary antibodies in 2% BSA. Primary antibodies used were: Paired-box 7 (PAX7, rabbit, 1:200, Abcam), anti-fibroblast antibody clone TE7 (mouse, 1:200, Millipore Sigma), MYOD (mouse, 1:40, Santa Cruz Biotechnology), myosin heavy chain (MyHC MF20, mouse, 1:100, DSHB), fatty acid binding protein 4 (FABP4, rabbit, 1:200, Abcam), fibronectin (FN, mouse, 1:200, Novus Biologicals), α-smooth muscle actin (α SMA, rabbit, 1:200) and platelet-derived growth factor α (PDGFRa/CD140a, goat, 1:200, R&D systems). The cells were then washed twice with 2% BSA and incubated with secondary antibodies in 2% BSA for 1 hour at room temperature. Donkey anti-mouse, donkey anti-rabbit and -rat, donkey anti-goat IgG conjugated to Alexa-Fluor (AF) 568, AF488, and AF647, respectively, were used as secondary antibodies with 1:500 dilutions (Life Technologies). Phalloidin (1:400, Invitrogen) and nuclear stain with DAPI (1:1,000, Invitrogen) were done in conjunction with the secondary antibodies. The cells were then washed twice and imaged on a Leica Thunder widefield microscope (Leica Microsystems). All images were analyzed using ImageJ software^9^.

### Immunofluorescence Image Analysis

Immunofluorescence images for SC and FAP lineage markers and multilineage differentiation were analyzed using custom macro scripts in ImageJ. Cell lineage was confirmed using colocalization analysis. Colocalization of DAPI^+^ nuclei, Pax7^+^, and MyoD^+^ features indicated SC lineage and colocalization DAPI^+^ nuclei and PDGFRα^+^ features indicated FAP lineage. Fusion index was quantified for SCs subjected to myogenic differentiation as the number of DAPI^+^ nuclei within MyHC^+^ myotubes divided by the total number of nuclei. To account for clusters of overlapping nuclei that could not be individually discriminated at day 2 and day 4 timepoints, the area of individual DAPI^+^ features was divided by the median nuclear area at day 0, then rounded up to the next whole number. FAP fibrogenic differentiation capacity was quantified as the mean fluorescence intensity (MFI) of FN and αSMA within PDGFRα^+^ segmented regions. FAP adipogenic differentiation capacity was quantified as total FABP4 MFI divided by the number of nuclei.

### Flow Cytometry

FAP and SC lineage was corroborated using flow cytometry. A complete flow cytometry protocol is included in Supplemental Protocol I. Briefly, FAPs and SCs were expanded in proliferation media to 80% and 70% confluency, respectively. Cells were then lifted using Tryp-LE (Gibco) and centrifuged at 400g for 5 minutes. The cell pellets were resuspended in warmed staining buffer (2.5% FBS in Dulbecco’s Phosphate Buffered Saline, DPBS) and counted. Cells were stained with anti-CD56-PE-CF594 (1:40, BD Bioscience), anti-CD31-APC-CY7 (1:40, BD Biosciences), anti-CD45-FITC (1:40, BD Biosciences), anti-CD140a/PDGFRa-PE (1:40, BD Biosciences), and Ghost Dye Violet 540 (1:100, Cytek) for 30 minutes at 4°C. A 5-laser Cytek Aurora with SpectroFlo software was used for quantification. UltraComp eBeads (Invitrogen) were used for compensation controls and spectral unmixing. Fluorescence minus one (FMO) controls were used to set gates for viability (Fig. S1A), CD45 (Fig. S1B), and CD31 (Fig. S1C). Unstained FAPs and SCs were used to set gates for CD56 and PDGFRα (Fig. S1D) due to insufficient numbers of cells in the CD56 and PDGFRα FMO controls (Fig. S1E-F). OMIQ software (Insightful Science) was used for flow cytometry data processing.

### 3D Tissue Engineered Muscle Constructs

Isolated SCs were incorporated into tissue engineered constructs to determine the capability of the cells to generate aligned bulk tissues. These muscle constructs were engineered using methods similar to those previously published^26,27^. A 24 well plate was prepared by coating wells with Silicone (Sylgard 184, Dow), positioning custom machined PolyEtherEther-Ketone (PEEK) micro-post inserts, and curing the silicone. Constructs were cultured on micro-posts to provide mechanical boundary conditions, anchor the constructs during hydrogel contraction, and facilitate myotube alignment. The plates were then sterilized by ethylene oxide before seeding cells. The constructs included hydrogel composed of type I collagen at a final concentration of 1.5 mg/mL (Advanced Biomatrix) and 20% Matrigel by volume (Corning). Prior to hydrogel polymerization, human SCs (passage 5) were resuspended in the collagen-Matrigel solution at a concentration of 4×10^6^ cells/mL and seeded onto each construct at 2×10^5^ cells per well. Constructs polymerized for 15 minutes at 37°C inside the incubator, and then 1.5 mL proliferation media was added to each well. Proliferation media was changed every 2 days in culture. After 4 days of proliferation, the media was switched to myogenic differentiation media. After 14 days of differentiation (18 total days of culture), the constructs were fixed in 4% PFA for immunofluorescence imaging and subsequent analysis for viability and differentiation. Viability was determined by staining with propidium iodide (Thermo Fisher Scientific), staining dead cells, and DAPI, staining total nuclei.

### Statistical Analysis

All statistical analyses were performed using GraphPad Prism 10. Data are presented as mean ± standard deviation. The statistical test and sample size used for each analysis are indicated in the relevant figure captions. Two-tailed unpaired t-tests were used for analyses comparing two groups across one variable. One-way analysis of variance (ANOVA) with Tukey’s multiple comparisons was used for analyses with one variable and more than two groups. Two-way ANOVA with Tukey’s or Dunnett’s multiple comparisons was used for analyses with two or more variables and two or more groups. The significance level (α) was set at p<0.05.

## Results

### FAPs and SCs retain expression of classical lineage-specific markers post-thaw

Lineage marker expression of previously harvested, sorted, and cryo-shipped FAPs and SCs^17^ was confirmed via IF imaging (Fig. 2A-C) and flow cytometry (Fig. 2D & E). For IF imaging analysis, SCs were stained for the canonical SC marker Paired box 7 (Pax7), Myoblast Determination Protein 1 (MyoD), filamentous actin (F-Actin), and the nuclear stain DAPI (Fig. 2A). FAPs were stained for the canonical FAP marker platelet-derived growth factor receptor α (PDGFRα), the anti-fibroblast clone TE7, and DAPI (Fig. 2B). An average of 91.6±1.4% of SCs were Pax7^+^MyoD^+^, while 7.3±1.4% of SCs were Pax7^+^MyoD^-^, and 0.58±0.077% were Pax7^-^MyoD^+^ (Fig. 2C; n=4 wells/cell type).

**Figure 2:**
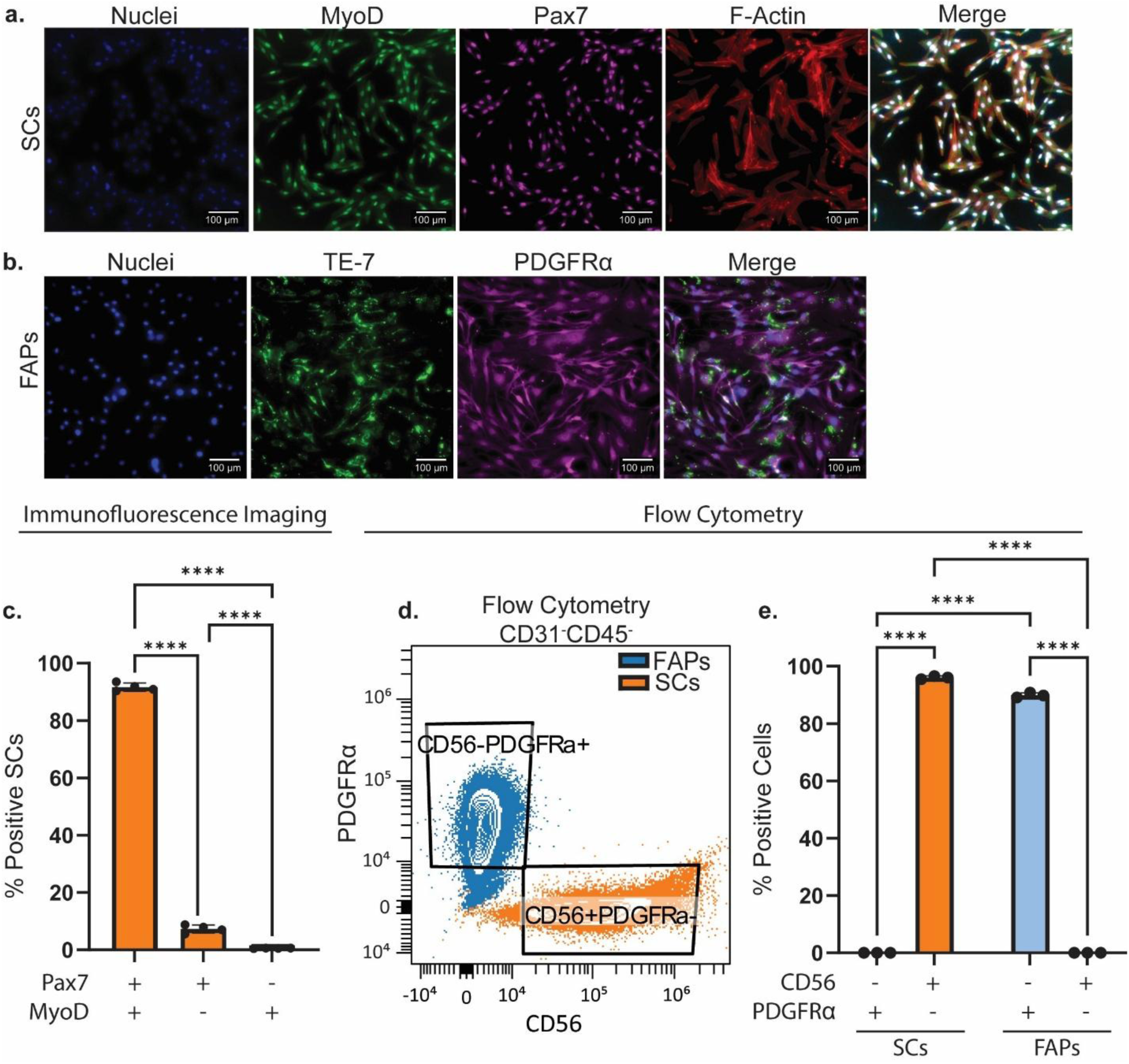
SCs and FAPs retain expression of classical lineage-specific markers post-thaw. A) Representative IF images of SCs after initial expansion post-thaw. SCs were stained for nuclei with DAPI (blue), for the activated SC marker MyoD (green), for the canonical SC marker Pax7 (purple), and the cytoskeletal protein F-Actin (red). B) Representative IF images of FAPs after initial expansion post-thaw. FAPs were stained for nuclei with DAPI (blue), for anti-fibroblast clone TE-7 (green), and the canonical FAP surface marker PDGFRα (purple). Scale bar = 100µm. C) IF images of SCs were quantified for the proportion of the population that stained for either Pax7 and MyoD (91.6±1.4%), Pax7 alone (7.3±1.4%), or MyoD alone (0.58±0.077%), shown as mean ± SD. n=4 wells/group, 5 replicate images per well. Statistics were one-way ANOVA with Tukey’s post-hoc test. ****p<0.0001. D) Flow cytometry of live CD31-CD45-cells from FAP samples (blue) and SC samples (orange) on a PDGFRα vs. CD56 biplot. E) Percentage of live cells that were either CD56-PDGFRα+ or CD56+PDGFRα-in SC samples (orange; 96.1±0.5% CD56+PDGFRα-) and FAP samples (blue; 89.9±0.9% CD56-PDGFRα+; n=3 technical replicates/sample type). Statistics were two-way ANOVA with Fisher’s post-hoc. ****p<0.0001.

We next performed flow cytometry to discriminate between FAPs and SCs. This panel consisted of CD56 to identify SCs, PDGFRα to identify FAPs, CD45 to exclude leukocytes, CD31 to exclude endothelial cells, and Ghost Dye to identify live cells. The gating scheme is illustrated in Supplemental Fig. 2. FAPs (blue) and SCs (orange) formed distinct clusters on a CD56 vs. PDGFRα biplot. An average of 96.1±0.5% of CD31^-^CD45^-^ cells in the SC samples were gated as CD56^+^PDGFRα^-^, and an average of 89.9±0.9% of CD31^-^CD45^-^ cells in the FAP samples were gated as CD56^-^PDGFRα^+^(Fig. 2E; n=3/group). Together, these IF and flow cytometry results confirm that both FAP and SC samples received from the HPA biobank are distinct populations of cells that maintain their lineage-specific markers post-thaw.

### Satellite cells maintain myogenic differentiation capacity after cryopreservation and shipping

Myogenic differentiation and fusion capacity of SCs is a crucial part of muscle regeneration after injury. After confirming SC expression of lineage markers against previously reported data^17^, we induced myogenic differentiation of SCs using Supplemental Protocol E. SCs were cultured in differentiation media for 0 (Fig. 3A), 2, (Fig. 3B), and 4 (Fig. 3C) days, after which they were fixed and stained for DAPI, MyHC, and F-Actin. We observed fusion of SCs into MyHC^+^ myotubes, with fusion indices of 0.3±0.16% at day 0, 10.3±1.8% at day 2 (p<0.0001 vs. day 0), and 25.4±2.1% at day 4 (p<0.0001 vs. day 0) (Fig. 3D). These results demonstrate preserved myogenic differentiation capacity of SCs following cryopreservation.

**Figure 3:**
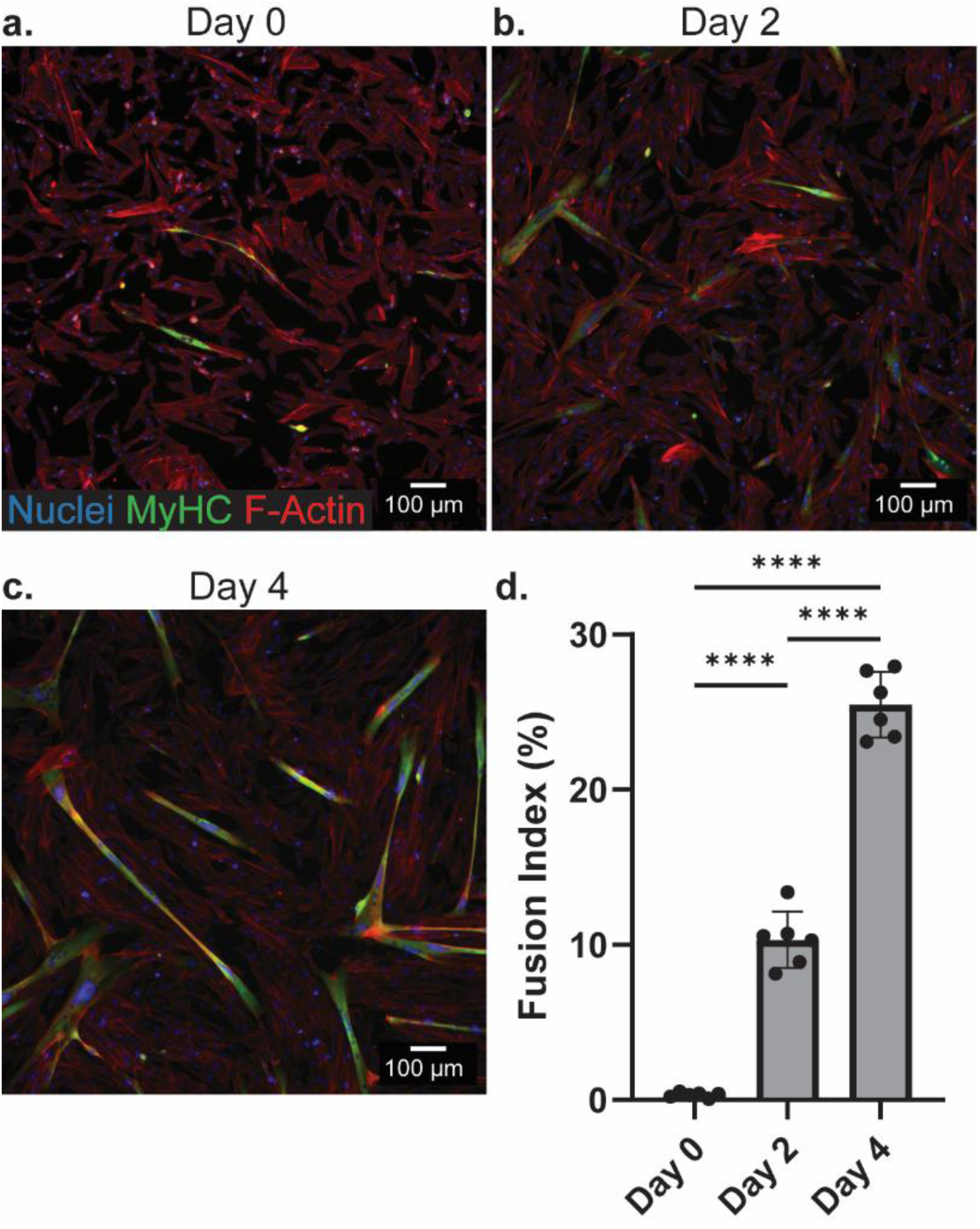
SCs retain myogenic differentiation capacity after biobanking. A-C) Representative IF images of SCs stained for nuclei (blue), myosin heavy chain (MyHC; green), and F-Actin (red) at day 0 (A), day 2 (B), and day 3 (C) of myogenic differentiation. Scale bar = 100µm. D) Fusion index was quantified at each timepoint (n=6 wells/timepoint; 0.3±0.16% at day 0, 10.3±1.8% at day 2, 25.4±2.1% at day 4). Data are shown as mean ± SD. Statistics were a one-way ANOVA with Tukey’s post-hoc. ****p<0.0001.

### FAPs maintain fibrogenic differentiation capacity after cryopreservation and shipping

Fibrogenic differentiation of FAPs into myofibroblasts is a hallmark pathological behavior of FAPs in conditions such as volumetric muscle loss. Thus, we assessed the fibrogenic differentiation capacity of human FAPs after cryopreservation. Fibrogenic differentiation *in vitro* is stimulated using TGF-β1. FAPs were cultured for 4 days in differentiation media containing either the proliferative growth factor bFGF alone (Fig 4A), bFGF and TGF-β1 (Fig. 4B), or TGF-β1 alone (Fig. 4C). Immunostaining was performed for αSMA, an intracellular marker of myofibroblasts and FN, a common extracellular matrix protein produced by FAPs, fibroblasts, and myofibroblasts. The mean fluorescence intensity (MFI) of both FN (Fig. 4D) and αSMA (Fig. 4E) were used to quantify fibrogenic differentiation of FAPs. TGFβ treatment with or without bFGF increased the MFI of FN compared to the bFGF-alone condition (p=0.0016 for bFGF vs. bFGF+TGFβ; p=0.0185 for bFGF vs TGFβ), with a similar trend for αSMA (p=0.0003 for bFGF vs. bFGF+TGFβ; p=0.0013 for bFGF vs TGFβ). These data indicate that FAPs can successfully undergo fibrogenic differentiation following cryopreservation.

**Figure 4:**
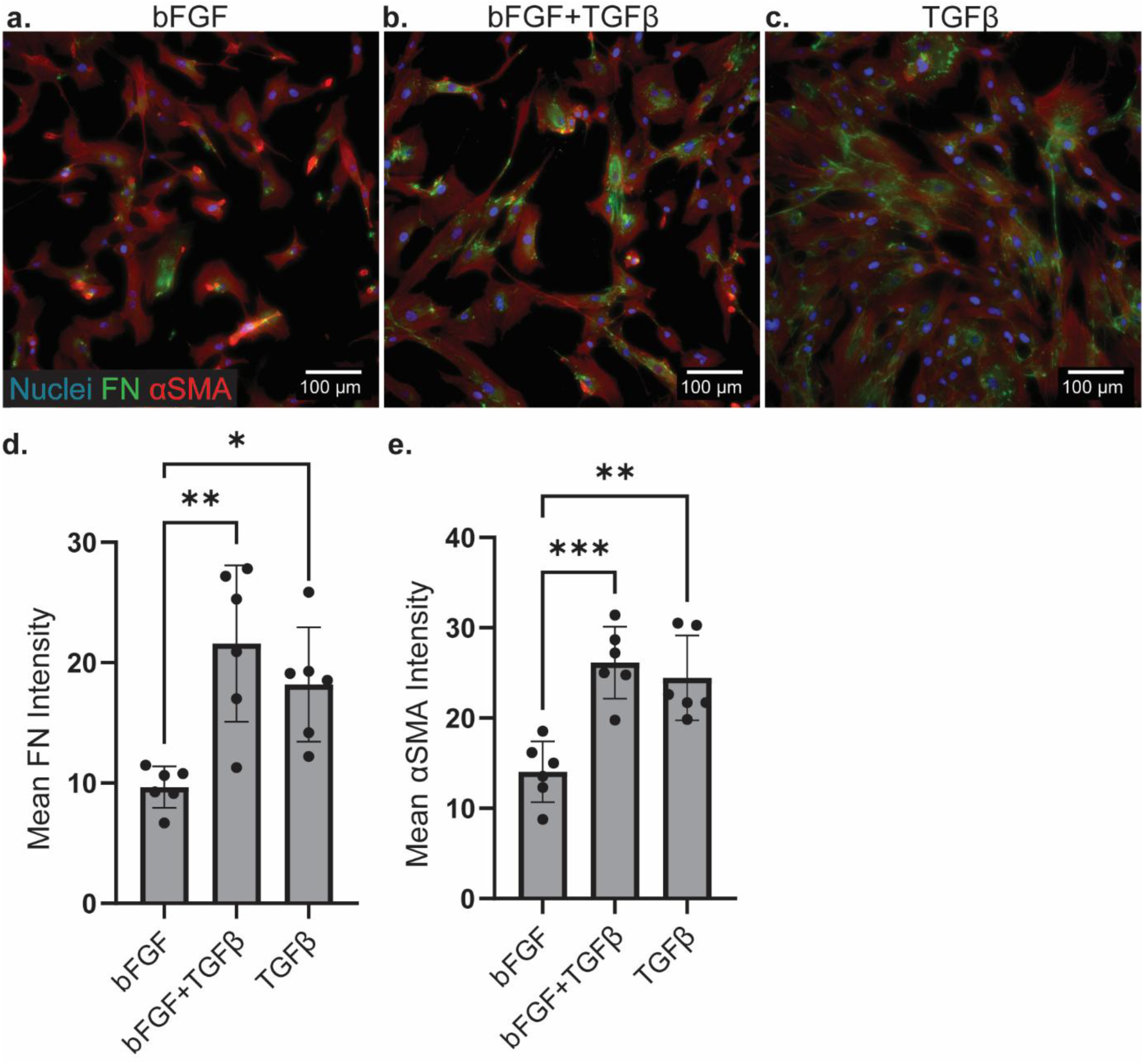
FAPs retain fibrogenic differentiation capacity after biobanking. A-C) Representative IF images of FAPs that were stained for nuclei (blue), FN (green), and αSMA (red) after culture with either bFGF alone (A), bFGF + TGFβ1 (B), or TGFβ1 alone (C). Scale bar = 100µm. D) Mean FN intensity within PDGFRα+-segmented regions was 9.66±1.72 for bFGF, 21.58±6.50 for bFGF+TGFβ1, and 18.19±4.75 for TGFβ1. E) Mean αSMA intensity within PDGFRα+-segmented regions was 14.06±.3.37 for bFGF, 26.15±3.98 for bFGF+TGFβ1, and 24.48±4.69 for TGFβ1. n=6 wells/group. Statistics were a one-way ANOVA with Tukey’s post-hoc. *p<0.05, **p<0.01, ***p<0.001.

### FAPs maintain adipogenic differentiation capacity after cryopreservation and shipping

To assess post-thaw adipogenic differentiation capacity, FAPs were cultured in either proliferation media or adipogenic differentiation media for 21 days. We then fixed and immunostained for FABP4, a protein associated with lipid droplets in adipocytes, F-Actin to visualize cell bodies, and DAPI for nuclei (Fig. 5A and B). Additionally, Oil Red O staining allowed for qualitative assessment of lipid droplet formation in FAPs cultured in proliferation media (Fig. 5C) versus those cultured in adipogenic differentiation media (Fig. 5D). Quantification of FABP4 MFI normalized to the number of nuclei (Fig. 5E) revealed increased expression of FABP4 in FAPs cultured in adipogenic differentiation media compared to FAPs cultured in proliferation media (p=0.0024, n=3 wells/group). Interestingly, the FABP4 staining observed in these images was predominantly localized to nuclei, rather than the cytoplasm. Collectively, these results support that FAPs retain adipogenic differentiation capacity following cryopreservation.

**Figure 5:**
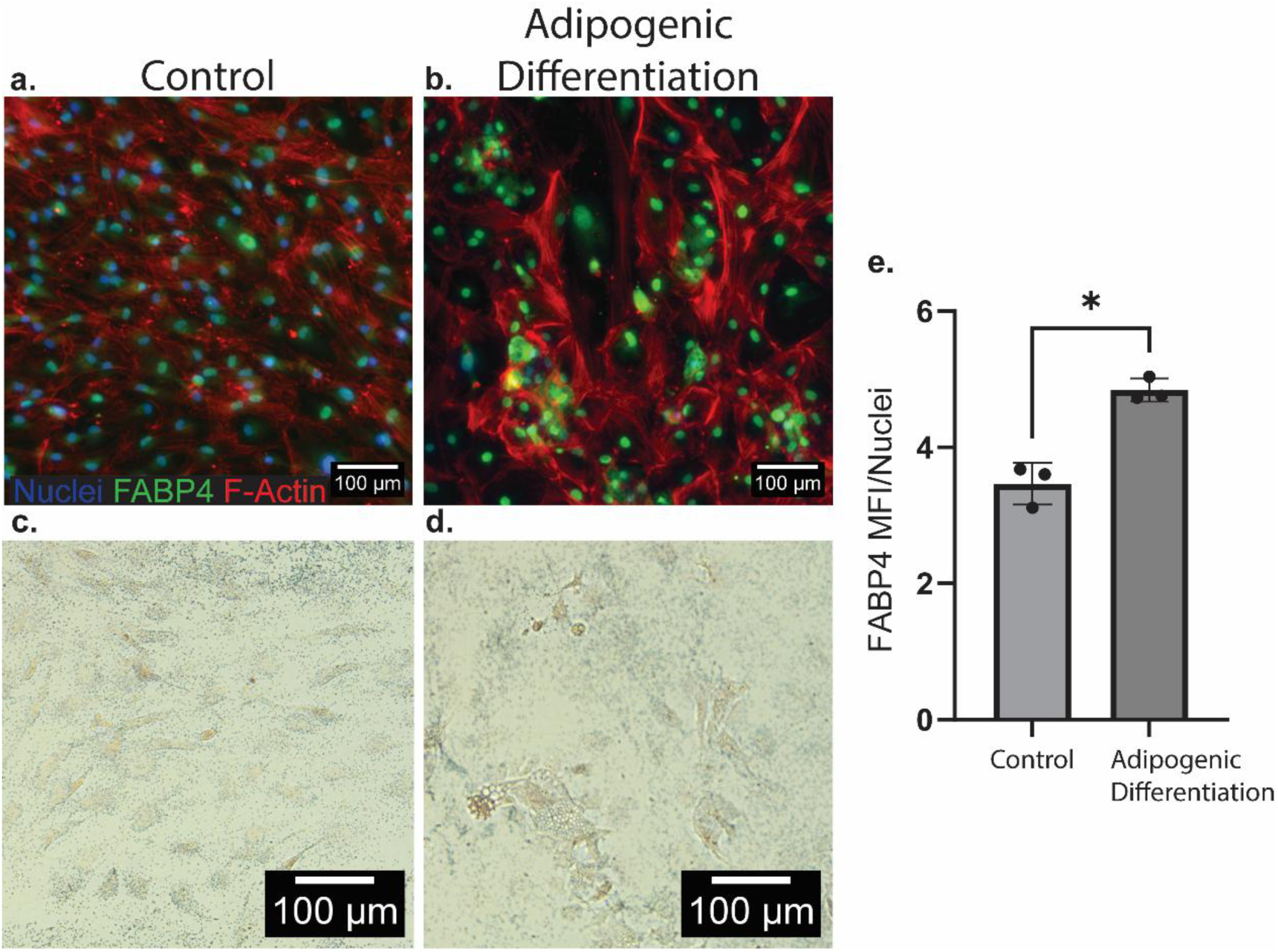
FAPs retain adipogenic differentiation capacity after biobanking. FAPs were cultured in either normal proliferation media (control; A, C) or adipogenic differentiation media (B, D), then stained for nuclei (blue), FABP4 (green), and F-actin (red) (A-B), or for lipid droplets using Oil Red O (C-D). Scale bar = 100µm. E) Mean FABP4 intensity per nuclei was 3.46±0.31 for control media and 4.84±0.18 for adipogenic differentiation media. n=3 wells/group. Statistics were a two-tailed unpaired t-test. *p<0.05.

### Previously biobanked satellite cells can be utilized in a 3D muscle organoid

To demonstrate the utility of primary human skeletal muscle cells obtained though this pipeline for use in NAMs, we incorporated SCs into a 3-dimensional muscle organoid. Brightfield images demonstrate visual formation of the SC-matrigel construct between day 0 (Figure 6A) and day 18 (Figure 6B) of culture, consistent with previous reports^26,27^. Furthermore, the differentiated myotubes within 3D muscle organoid exhibited aligned myotubes (Figure 6C) with high viability (73±16% live cells). These 3D muscle organoids successfully recapitulate the anisotropic and highly organized architecture of skeletal muscle, supporting the use of previously biobanked cells to generate human-specific skeletal muscle NAMs.

**Figure 6:**
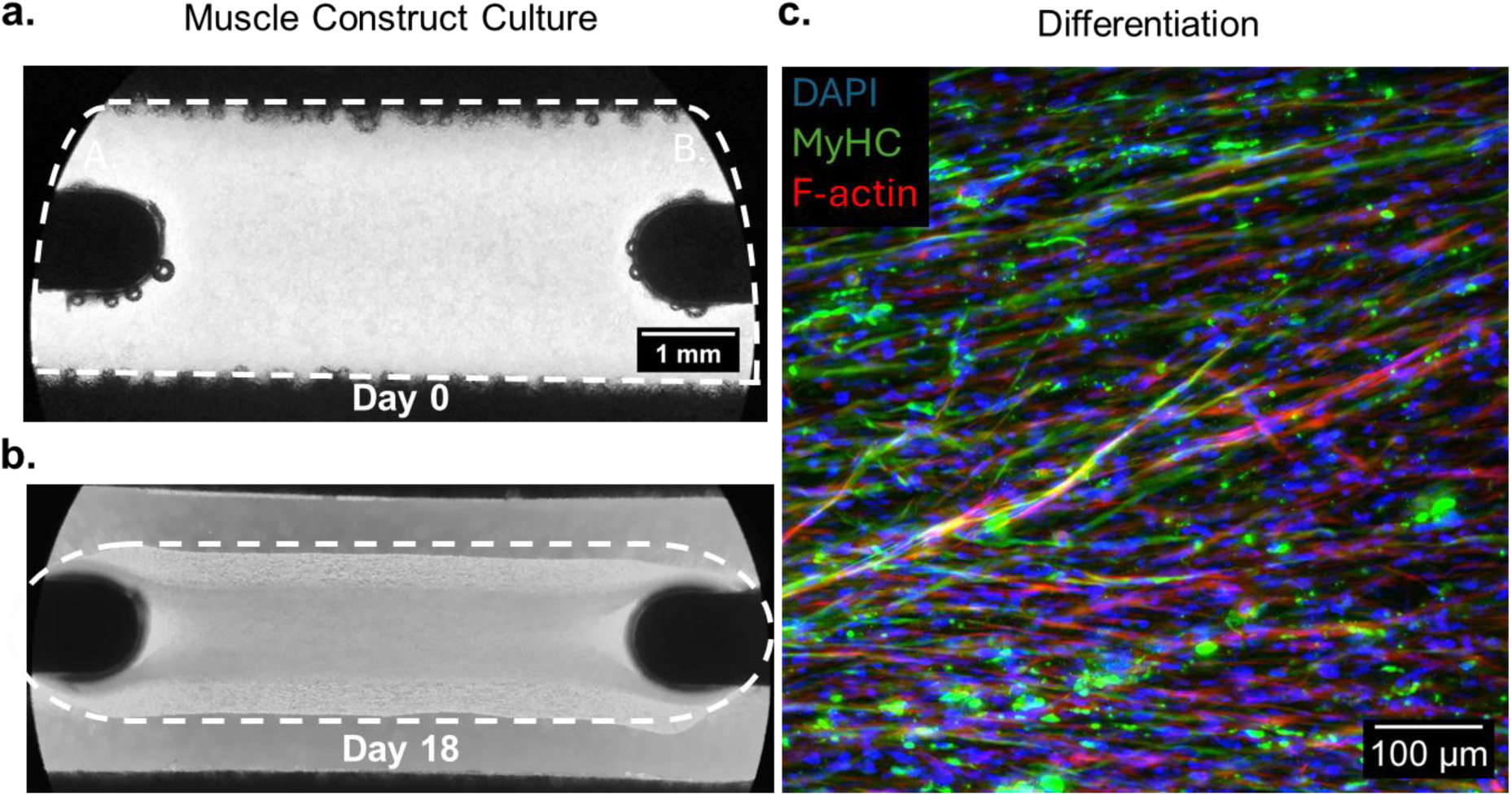
Biobanked and cryo-shipped human SCs were used to generate 3D tissue-engineered muscle constructs. SCs seeded on micro-posts on day 0 (A) showed definitive formation of a construct by day 18 of culture (B). C) Immunofluorescence imaging confirmed that SCs within the constructs had differentiated into aligned, multinucleated myotubes. Differentiated myotubes within the 3D muscle construct were stained for nuclei (DAPI, blue), myosin heavy chain (MyHC; green), and F-Actin (red).

## Discussion

This study addresses critical gaps in current workflows using FAPs and SCs. We leverage foundational co-isolation protocols^17^ to establish an end-to-end pipeline for biobanking, shipping, and utilizing these donor-matched cell populations.

Previously biobanked SCs and FAPs from one young adult donor were thawed, expanded, passaged, and cryopreserved. Immunofluorescence imaging supported that expanded SCs express Pax7 and MyoD, while FAPs express PDGFRα/CD140a and TE7. Importantly, we confirmed the retention of lineage specific markers in these previously biobanked SC and FAP samples using flow cytometry. The biplot of CD56 vs. PDGFRα generated from this flow cytometry analysis showed minimal overlap of SC and FAP samples, indicating that they are distinct cell populations. We then confirmed that previously biobanked SCs maintain robust capacity for myogenic differentiation, with day 4 fusion indices in line with previous reports^17^ (Fig. 3). We also report retention of fibrogenic (Fig. 4) and adipogenic (Fig. 5) differentiation capability of post-thaw FAPs. Lastly, we demonstrate the potential utility of these cells for the NIH’s recent initiative for NAMs by incorporating SCs in a 3D tissue-engineered muscle construct (Fig. 6).

The verification experiments performed in this study, using cells from a single donor, demonstrate a critical proof-of-concept of whether the biobanking pipeline can preserve the ontological identity and differentiation capacity of FAPs and SCs. The present results were somewhat limited by the collection of images in a single plane, rather than a z-stack. Specifically, signal intensity was quantified at the nuclear plane, rather than throughout the 3-dimensional cell structure. Although this approach precludes cell-wide assessment of signal intensity, comparisons across planar views support that the differentiation capacity was preserved in these MuSCs and FAPs following biobanking.

While this study primarily leveraged immunofluorescence techniques to assess changes in cellular architecture before and after differentiation, further studies are needed to characterize cellular function after biobanking, including intercellular signaling, metabolism, and multi-omic signatures. Furthermore, the assessment of donor-to-donor variability, a known challenge in primary human cell research, was beyond the scope of the present study. Future efforts should leverage this established pipeline to investigate the variability of cellular function between donors. Expansion of the Biobank to include donor samples of diverse age, sex, ethnicity, disease, and metabolic characteristics will further enable investigations into how these patient-specific variables influence SC and FAP behavior. The present assessment lacks a direct comparison of pre-freeze/post-freeze cellular characteristics for the same cell populations, as cells were biobanked at the isolation site and utilized at an independent facility. However, this pipeline reflects real-world biobank usage and demonstrates the utilization of post-thaw SCs and FAPs upon receipt.

This pipeline will directly benefit research pursuits in regenerative medicine that were previously constrained by the limited availability of standardized primary human muscle cells. For example, the demonstrated utility of post-thaw SCs for generating a 3D tissue-engineered muscle construct positions this biobanking pipeline to accelerate the development of clinically relevant, human-based NAMs for disease modeling and therapeutic screening. Because cells from the same donor share a genome, disease state, and metabolic history, the use of donor-matched SC-FAP pairs will enable patient-specific NAMs to study the multi-cellular mechanisms which drive pathological regeneration in clinical contexts including volumetric muscle loss. Finally, the standardized biobanking infrastructure established by this work will support multi-site reproducibility and cross-laboratory comparisons on cells from the same donor sample, amplifying the effect skeletal muscle cellular studies on the field of regenerative medicine.

## Data Availability

The data and ImageJ macro code are available from the corresponding author upon reasonable request.

## Acknowledgements

Custom-machined micro-post inserts used in the 3D muscle organoid were created by the Oregon Fabrication and Design (OFAD) group. Figure 1 was created in BioRender. Willett, N. (2026) https://BioRender.com/hupwot8.

## Funding

This work was supported by the Wu Tsai Human Performance Alliance and the Joe and Clara Tsai Foundation, as well as support from the National Institutes of Health (T32GM008666 to A.B.; 3R01AR078375-05S1 to N.J.W.), the Rita L. Atkinson Graduate Fellowship (to A.B.) and Cheng Family Innovative Research Award (to A.B.).

## Competing Interests

The authors declare no competing interests.

## Author Contributions

F.S.P., A.R., G.E.P., A.B., S.R., A.J.E., S.R.W., and N.J.W. conceived and designed the research. C.M.R. harvested tissue samples. F.S.P, A.R., G.E.P., A.B., and S.R. conducted the experiments. F.S.P., A.R., G.E.P., A.B., S.R., S.R.W., and N.J.W. analyzed and interpreted the results. F.S.P., G.E.P., and A.R. prepared figures. F.S.P., A.R., G.E.P., A.B., S.R., S.R.W., and N.J.W. drafted the manuscript. F.S.P., A.R., G.E.P., A.B., S.R., R.E.G., A.J.E., S.R.W., and N.J.W. edited and revised the manuscript. A.B., R.E.G., A.J.E, S.R.W., and N.J.W. acquired funding. All authors approved final version of the manuscript.

## Notes

### Competing Interest Statement

The authors have declared no competing interest.

