## Supplementary Methods and Figures for "A Multi-Institution Biobanking Pipeline for Primary Human Satellite Cells and Fibro-Adipogenic Progenitors"

**Materials:**

- Biosafety Cabinet
- Water bath (37°C)
- Incubator (37°C)
- Treated culture flask (Thermo Fisher Scientific, Cat# 159910)
- Well plates (CellTreat Scientific, Cat# 229123 and Cat# 229112)
- Centrifuge
- Conical tubes
- Serological pipettes
- Pipette tips

- DMEM (Thermo Fisher Scientific, Cat# 11054020)
- Pen/Strep
- Fetal Bovine Serum (FBS)
- bFGF
- Stem Pro Adipogenesis differentiation kit (Gibco)
- DMSO
- -80°C freezer
- Liquid nitrogen
- Countess cell counter
- TrypLE
- Trypan blue

**Media Composition:**

- Basal Media:
  - DMEM (low glucose, - phenol red)
  - 1% P/S
  - 20% FBS
  - 1% P/S
- Growth Media:
  - Basal media
  - 5 ng/mL bFGF
- Adipogenic Differentiation Media:
  - Stem Pro Adipogenesis differentiation kit (Gibco A1007001)
- Fibrogenic Differentiation Media:
  - Basal media + 10 ng/mL TGFβ-1
- Myogenic Differentiation Media:
  - DMEM (low glucose, - phenol red)
  - 5% Horse Serum
  - 10 µg/mL insulin
  - 1% P/S

**Supplemental Protocols:**

<u>A: Thawing and plating frozen cells (SCs and FAPs)</u>

1. Warm pre-mixed growth media in water bath until media has equilibrated at 37°C
2. Add 5mL of fresh prewarmed media to a 15mL falcon tube
3. Remove cells from frozen storage and immediately place in 37°C water bath for 2 minutes to thaw
4. Use a P1000 micropipette to carefully transfer cells to the falcon tube containing prewarmed growth medium. Dispense cells dropwise to minimize shear forces.
5. Centrifuge at 400xg for 5 mins at room temperature (RT)
6. Check pellet, spin ~ 3 minutes longer if pellet not well defined
7. Aspirate the supernatant and resuspend cells in 1mL of fresh pre-warmed media.
8. Count cells: 10 µL cell suspension + 10 µL trypan blue for Countess, record % live cells and number live cells/mL
9. Plate cells in flask (approximately 1 million cells in 27 mL prewarmed media for a T175 flask)
10. Add bFGF (target final concentration = 5 ng/mL) to flask
11. Incubate cells at 37°C/5% $CO_2$/~21% $O_2$

12. Replace growth medium 24 hours post plating with fresh growth media
    a. This step removes any remaining DMSO (present in storage solution, cytotoxic)
13. Record notes, including: the name of the cell line, the media components (and date mixed), media changes, calculation of the population doubling level (PDL), and any observations relative to morphology

<u>B: Media Change – Expansion (SCs and FAPs)</u>

1. Performed every 2-3 days during expansion
2. Warm growth media in water or bead bath until properly equilibrated to 37°C
3. Remove flask from incubator
4. Aspirate media from culture vessel
5. Add fresh prewarmed growth media to culture vessel (T175: 27 mL media)
6. Add bFGF (target final concentration = 5 ng/mL)

<u>C: Passaging</u>

1. Passage FAPs at 80-85% confluency. Passage SCs at 60-70% confluency.
2. Aspirate media from culture vessel
3. Wash the monolayer in DPBS (-CaCl -MgCl) that has been pre-warmed to 37°C
4. Gently agitate culture vessel containing DPBS
5. Aspirate DPBS
6. Add TrypLE to culture vessel (17 mL for T175)
7. Rotate the flask to completely cover the monolayer of cells
8. Incubate until cells detach from the surface (check under microscope after 3 minutes)
9. Neutralize trypLE with equal volume of media (17 mL for T175)
10. Transfer cell suspension to a labeled conical tube
11. Centrifuge (400 rcf, 5 min., room temperature)
12. Aspirate the supernatant and resuspend with 1 mL fresh prewarmed medium
14. Count the cells: 10 µL cell suspension + 10 µL trypan blue for Countess, record % live cells and number live cells/mL
13. Transfer the appropriate number of cells to a new labeled tissue culture vessel containing pre-warmed growth media

<u>D: Freezing cells</u>

1. Lift cells and count (see section C above)
2. Calculate the volume of cell suspension needed to store aliquots of 1 million cells
3. Pre-label appropriate number of cryovials
4. Aliquot vials
    a. V cell suspension
    b. (900 - V cell suspension) µL growth media
    c. 100 µL DMSO

i. ** Cells should be exposed to DMSO for as little time as possible prior to freezing, as it is cytotoxic

5. Place vials in a room-temperature Mr. Frosty container (with sufficient isopropanol)
6. Place the Mr. Frosty container in the -80℃ freezer overnight
7. After 24 hours, transfer to vials to long-term liquid nitrogen storage

E: Myogenic Differentiation (SCs)

1. Seed SCs at 20,000 cells/$cm^2$ in a 24-well plate
2. Culture cells in growth media until 100% confluency is reached, replacing growth media every 2 days
3. Once cells are ready, wash cells with 1x DPBS and add myogenic differentiation media composed of DMEM low-glucose with 5% horse serum, 10µg/mL insulin, and 1% P/S.
4. Replace differentiation media every 2 days
5. On day 4 of differentiation, use cells for immunofluorescence imaging (fix with 3.7% PFA)

F: Fibrogenic differentiation (FAPs)

1. Seed FAPs at 5,000 cells/$cm^2$ in a 24 well plate
2. Expand in growth media 24 hrs
3. Aspirate growth media and rinse cells with DPBS
4. Aspirate DPBS
5. Differentiate FAPs in 3 groups:
    a. Growth media (i.e. basal media + 5 ng/mL bFGF)
    b. Growth media + 10 ng/mL TGF-β1
    c. Basal media + 10 ng/mL TGF-β1
6. Replace differentiation media (groups a-c above) every day
7. On day 3 of differentiation, use cells for immunofluorescence staining (fix with 3.7% PFA)

G: Adipogenic differentiation (FAPs)

1. Seed FAPs at 10,000 cells/$cm^2$ in 12-well plate
2. Expand in growth media 4 days
3. Aspirate growth media and rinse cells with DPBS
4. Aspirate DPBS
5. Add adipogenic differentiation media (Stem Pro Adipogenesis Differentiation Kit, Gibco) and differentiate 14 days
    a. Replenish media every 3-4 days
6. On day 14 of differentiation, use cells for gene expression analysis (lyse cells for RNA collection) or immunofluorescence staining (fix with 4% PFA)

H: Immunofluorescence Staining (SCs and FAPs)

*DAY 1 – IF staining for SCs and FAPs*

1. Prepare Block Buffer: 2% BSA in PBS*
    a. Make enough for 2.50 mL/well (It will be used several times)
    b. Keep on ice/at 4°C for storage
    c. Formula
        i. BSA = “%” grams/100mL of water
        ii. E.g for 20 wells, prep at least 50 ml of Block buffer
            1. Weigh 1 g of BSA on microscale
            2. Measure PBS using serological pipette
            3. Flip up and down until solution is homogenous
2. Prepare Permeabilization Buffer: 0.2% Triton X-100 in PBS*
    a. E.g. for 20 wells of 24 well plates, prepare 10 ml of Perm Buffer
        i. 10ml PBS + 20 uL of Triton-X-100
        ii. Flip up and down until solution is mixed (may take 5-10 minutes to mix)
3. Aspirate media and wash cells with DPBS
    a. 500 µL 1X DPBS per well in a 24-well plate
4. Fix Cells
    a. Aspirate DPBS
    b. Add enough 4% PFA to cover each well – 500 µL
    c. Incubate at RT for 10-15 minutes
    d. Wash each well twice with 1X DPBS
5. Permeabilize Cells:
    a. Add 500 µL Permeabilization Buffer
    b. Incubate at RT for 10 minutes
    c. Wash twice with blocking buffer for 5 minutes at a time
6. Block:
    a. Add 500 µL Blocking Buffer
    b. Incubate for 1 hr at RT
7. While cells are incubating in block buffer, prepare 1° antibody staining solution with 2% BSA in DPBS and the following dilution factors for each antibody.
    a. For FAPs lineage markers
        i. Anti-PDGFRa – 1:200 (Goat host, R&D systems AF-307-NA)
        ii. Anti-TE7 – 1:200 (Mouse host, Millipore Sigma CBL271)
    b. For SC lineage markers
        i. Anti-Pax7 – 1:200 (Rabbit host, Abcam ab187339)
        ii. Anti-MyoD – 1:40 (Mouse host, Santa Cruz PA5-30591)
    c. For FAPs fibrogenic differentiation
        i. Anti-FN – 1:200 (Mouse host, Novus Biologicals NBP1-51723)
        ii. Anti-aSMA – 1:200 (Host Rabbit, Abcam ab124964)
        iii. Anti-PDGFRa – 1:200 (Goat host, R&D systems AF-307-NA)
    d. For FAPs adipogenic differentiation
        i. Anti-FABP4 – 1:200 (Rabbit host, Invitrogen PA5-30591)
    e. For SC myogenic differentiation
        i. Anti-MHC-I – 1:100 (Mouse host, DSHB MF-20)
8. Stain with 1° Antibody:
    a. Aspirate Blocking Buffer – No need to wash

b. Add 350 uL staining solution to each well (directions for 24 wellplate. Adjust for other sizes)
c. Wrap plate in parafilm to prevent evaporation
d. Incubate at 4°C overnight (16-18 hours)

*DAY 2 – IF Staining for SCs and FAPs*

9. Prepare the following 2° antibody staining cocktails in 0.2% BSA. All staining cocktails have 300 nM DAPI and a 1:500 dilution of each 2° antibody. Mix gently without vortexing, and protect from light.
    a. For FAPs lineage markers
        i. Donkey anti-Goat IgG AF647 (Invitrogen A21447)
        ii. Donkey anti-Mouse IgG AF488 (Invitrogen A21202)
    b. For SC lineage markers
        i. Donkey anti-Rabbit IgG AF568 (Invitrogen A10042)
        ii. Donkey anti-Mouse IgG AF488 (Invitrogen A21202)
        iii. Phalloidin AF568 – 1:400 (Thermo Scientific A12380)
    c. For FAPs fibrogenic differentiation
        i. Donkey anti-Mouse IgG AF488 (Invitrogen A21202)
        ii. Donkey anti-Rabbit IgG AF568 (Invitrogen A10042)
        iii. Donkey anti-Goat IgG AF647 (Invitrogen A21447)
    d. For FAPs adipogenic differentiation
        i. Donkey anti-Rabbit IgG AF488 (Invitrogen A21206)
        ii. Phalloidin AF568 – 1:400 (Thermo Scientific A12380)
    e. For SC myogenic differentiation
        i. Donkey anti-Mouse IgG AF488 (Invitrogen A21202)
        ii. Phalloidin AF568 – 1:400 (Thermo Scientific A12380)
10. Stain with 2° Antibody Staining Solution
    a. Wash each well twice with blocking buffer
    b. Add 350 μL of 2° Antibody Staining Solution to each well (for 24 well plates)
    c. Cover plate with foil (Make sure there are no holes or tears in the foil to keep the fluorescent dyes protected from light)
    d. Incubate at RT for 30 minutes
    e. Wash 2 times with 1X PBS
11. Optional: use mounting medium Fluoromount-G (SouthernBiotech)
    a. Aspirate all liquid
    b. Place "one drop" in each well
    c. Lift well walls
    d. Place cover slip
12. Visualize under a microscope with the appropriate fluorescent filters
    a. DAPI excitation/emission (ex/em): I = 360nm/465nm
    b. Alexafluor 488 ex/em: I = 490nm/520nm
    c. AlexaFluor 568 ex/em: I = 578nm/603nm
    d. Alexafluor 647 ex/em: I = 650nm/665nm

I: Flow Cytometry Analysis of SCs and FAPs

1. Lifting cultured cells (In BSC)
    a. After expanding SCs and/or FAPs in a T175 flask, aspirate cell culture media
    b. Wash with 25mL 1X DPBS and aspirate
    c. Lift cells from flask using 10 mL 0.25% Trypsin-EDTA for 5 minutes.
    d. After 5 minutes, check for cell adhesion under brightfield microscope. Tap bottom of flask if cells are still adherent.
    e. Dilute Trypsin-EDTA with an equal volume (10mL) of cell culture media.
    f. Strain cells through a 40um cell strainer into a fresh 50 mL conical tube. Rinse cell strainer with 10mL of media.
    g. Centrifuge cells at 400g for 5 minutes.
    h. Aspirate supernatant and resuspend cell pellet in 1mL warmed staining buffer
    i. Take a 10uL sample of the cell suspension and mix with an equal volume of Trypan Blue for cell counting on a Countess.
2. Split cells, viability staining, and fixation (In BSC)
    a. Transfer cells to their respective FACS tubes for staining.
        i. Between 100,000-250,000 cells per well for each unstained control, FMO control and viability single stain control
        ii. >250,000 cells per well for each fully-stained sample.
    b. Equalize the volume of each tube that contains cells. Place a FACS cap on each tube. Centrifuge at 400g for 5 mins.
    c. During the centrifugation, prepare single-stain controls for compensation using beads:
        i. Add 300uL of DPBS into the 5 FACS tubes for:
            1. CD31 single-stain
            2. CD45 single-stain
            3. CD56 single-stain
            4. CD140a single-stain
            5. CD140b single-stain
        ii. Add one drop of UltraComp eBeads to each of these tubes.
    d. Once the centrifugation is done, carefully aspirate the supernatant without disturbing the pellet.
        i. Resuspend the unstained samples with 100uL of staining buffer. Set these tubes aside on ice to prevent accidentally staining with antibodies.
        ii. Resuspend fully-stained samples and FMOs in 99uL of stain buffer.
    e. Add 1uL of Ghost Dye Violet 540 to each fully-stained sample and FMO and gently mix. Do not add Ghost Dye Violet 540 to the viability FMO.
    f. Incubate Samples on ice for 30 minutes in the dark.
    g. After 30 minutes, add 1mL of DPBS to dilute the viability dye, then centrifuge at 400g for 5 minutes.
    h. Carefully aspirate the supernatant, and resuspend in 1mL of DPBS again.
    i. Retrieve the unstained samples you set aside, and add 1 mL of DPBS to those tube.

j. Centrifuge all tubes at 400g for 5 minutes.
k. Carefully aspirate the supernatant, and resuspend pellet in 100uL of 4% PFA. Incubate at room temperature for 15 minutes
l. After 15 minutes, add 1mL DPBS to dilute the PFA. Centrifuge at 400g for 5 minutes.
m. Carefully aspirate the supernatant. Resuspend the pellet in 1 mL of DPBS to wash. Centrifuge again at 400g for 5 minutes.

3. Antibody staining beads and fixed cells
   a. Carefully aspirate the supernatant without disturbing the pellet.
      i. Resuspend the FMO controls and fully-stained samples with the corresponding volume in the "**Staining media volume to add**" column of Table 1.
      ii. Set aside the unstained controls on ice to avoid accidentally staining them.
   b. Add the corresponding volume of the correct antibodies and to each FMO control tube as listed in the **"1º Ab volume"** column of Table 1.
      i. **NOTE: Do not add anti-CD56 to the CD56 FMO, anti-CD31 to the CD31 FMO, anti-CD45 to the CD45 FMO, anti-CD140a to the CD140a FMO, anti-CD140b to the CD140b FMO, or the viability stain to the viability FMO**
   c. Add the corresponding volume of the viability stain and all antibodies to the fully-stained samples.
   d. Incubate cells and beads for 30 minutes on ice in the dark.
   e. After staining, add 1mL DPBS to all tubes (including unstained) to wash. Replace caps on tubes and centrifuge at 400g for 5 minutes.
   f. Carefully aspirate supernatant without disturbing cell/bead pellet.
   g. Repeat wash step: resuspend pellets in 1mL of DPBS, replace caps, and centrifuge at 400g for 5 minutes.
   h. Carefully aspirate supernatant. Resuspend cells and beads in 200uL FACS buffer. Store at 4 degrees C until analysis on the flow cytometer.

| Fluorophore | Sample/Control Type | Tube Contents | 1º Ab volume (ratio; µL) | Staining media volume to add (µL) |
|---|---|---|---|---|
| Unstained | Unstained SCs | Live SCs | None | 100 uL |
| Unstained | Unstained FAPs | Live FAPs | None | 100uL |
| PE-CF594 (CD56) | Single-stain | Beads | 1uL | 300 µL |
| APC-Cy7 (CD31) | Single-stain | Beads | 1uL | 300 µL |
| FITC (CD45) | Single-stain | Beads | 1uL | 300 µL |
| PE (PDGFRa) | Single-stain | Beads | 1uL | 300 µL |
| Ghost Dye Violet 540 (viability) | Viability Single Stain | Live FAPs or SCs | 1uL | 99 µL |
| PE-CF594 (CD56) | FMO | Live SCs | 1uL viability 2.5uL per Ab | 89 µL |
| APC-Cy7 (CD31) | FMO | Live SCs | 1uL viability | 89 µL |

| | | | | |
|---|---|---|---|---|
| | | | 2.5uL per Ab | |
| FITC (CD45) | FMO | Live SCs | 1uL viability 2.5uL per Ab | 89 µL |
| PE (PDGFRa) | FMO | Live FAPs | 1uL viability 2.5uL per Ab | 89 µL |
| Ghost Dye Violet 540 (viability) | FMO | Live FAPs | 2.5uL per Ab | 87.5 µL |
| All | Fully stained | Live FAPs | 1uL viability 2.5uL per Ab | 86.5 µL |
| All | Fully stained | Live FAPs | 1uL viability 2.5uL per Ab | 86.5 µL |
| All | Fully stained | Live FAPs | 1uL viability 2.5uL per Ab | 86.5 µL |
| All | Fully stained | Live SCs | 1uL viability 2.5uL per Ab | 86.5 µL |
| All | Fully stained | Live SCs | 1uL viability 2.5uL per Ab | 86.5 µL |
| All | Fully stained | Live SCs | 1uL viability 2.5uL per Ab | 86.5 µL |

**Supplemental Figures:**

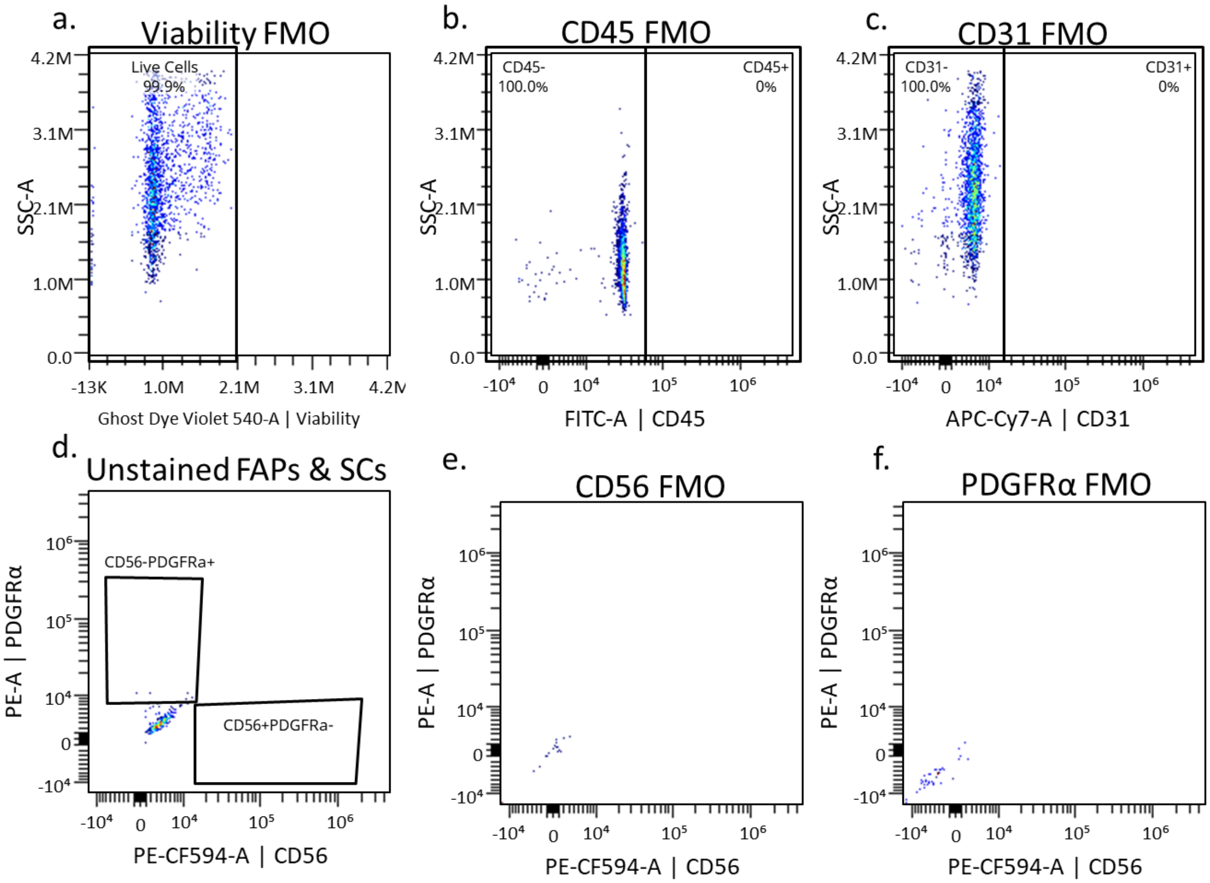


***Supplemental Figure 1: Flow cytometry gates were set on FMO controls and unstained cells.*** *A) The viability gate was set using an FMO control for Ghost Dye Violet 540. B) The CD45 gate was set on live cells at 5x10$^4$ using an FMO control for FITC. C) The CD31 gate was set on live cells at ~0.75x10$^4$ using an FMO control for APC-Cy7. D) The gates for CD56 (~0.5x10$^4$) and PDGFRα (1x10$^4$) were set using unstained FAPs an SCs. The CD56 (E) and PDGFRα (F) FMOs were not used to set gates due to an insufficient number of cells.*

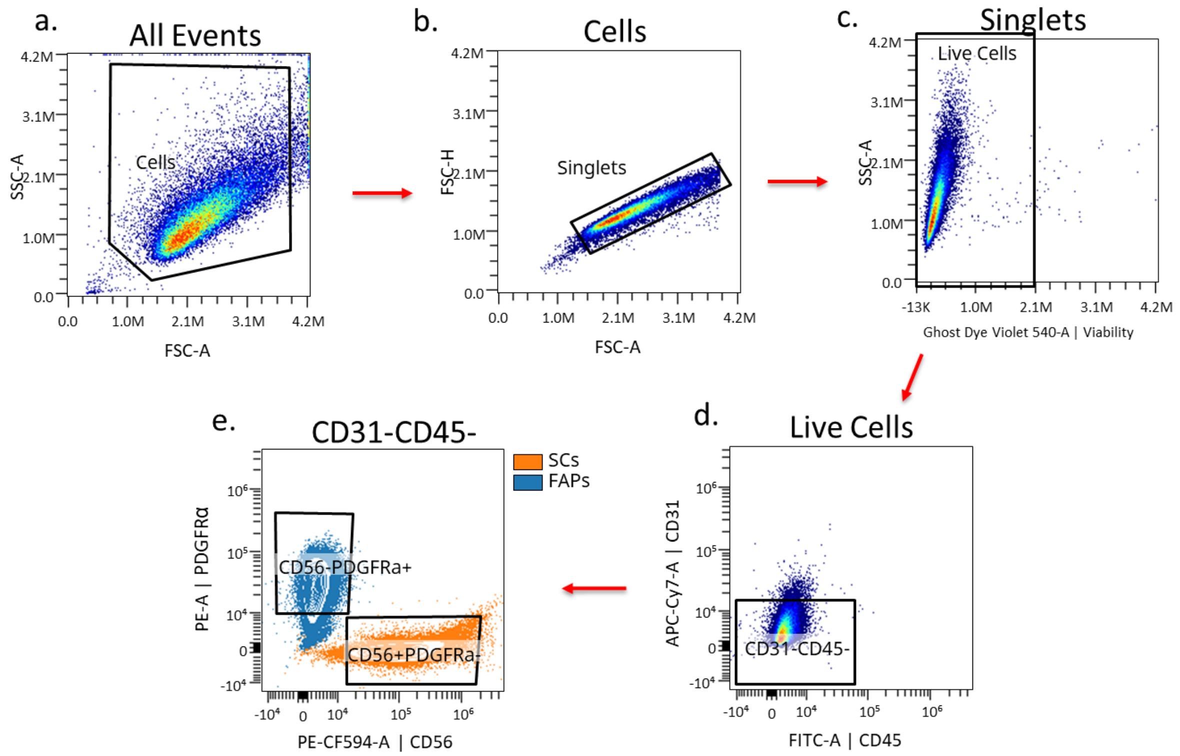


***Supplemental Figure 2: Representative flow cytometry gating scheme for SCs and FAPs.*** *A) Cells were selected from all events on FSC-A vs. SSC-A. B) Doublets were then excluded using FSC-A vs. FSC-H. C) Dead cells were then excluded based on their positivity for Ghost Dye Violet 540. D) Leukocytes (CD45+) and endothelial cells (CD31+) were excluded. E) CD56-PDGFRα+ FAPs (blue) and CD56+PDGFRα- SCs (orange) were selected for quantification. Contours represent cell density. n=3 technical replicates per cell type.*
